# Dynamic expression of MMP28 during cranial morphogenesis

**DOI:** 10.1101/850834

**Authors:** Nadege Gouignard, Eric Theveneau, Jean-Pierre Saint-Jeannet

**Affiliations:** Centre de Biologie du Développement (CBD), Centre de Biologie Intégrative (CBI), Université de Toulouse, CNRS, UPS, France; Department of Basic Science and Craniofacial Biology, New York University College of Dentistry, New York NY 10010, USA

**Keywords:** MMP, embryogenesis, cranial placodes, neural crest, Xenopus

## Abstract

Matrix metalloproteinases (MMP) are a large family of proteases comprising 24 members in vertebrates. They are well known for their extracellular matrix remodelling activity. MMP28 is the last member of the family to be discovered. It is a secreted MMP involved in wound healing, immune system maturation, cell survival and migration. MMP28 is also expressed during embryogenesis in human and mouse. Here we describe the detailed expression profile of MMP28 in *Xenopus laevis* embryos. We show that MMP28 is expressed maternally and accumulates at neurula and tailbud stages specifically in the cranial placode territories adjacent to migrating neural crest cells. As a secreted MMP, MMP28 may be required in normal neural crest-placode interactions.

## Background

Matrix metalloproteinases form a large family of proteases regrouping 23 members in human and 24 in *Xenopus laevis*^1^. MMPs are translated as zymogenes, or pro-enzymes, where the N-terminal part of the protein, called pro-domain, blocks the catalytic domain. This pro-domain can either change conformation (a process known as allosteric activation) or be removed by proteolytic cleavage to ensure the activity of the protein^2^. MMPs are Zn^2+^-dependent proteases mostly known for their extracellular matrix remodelling activity but they are also capable of cleaving a wide range of substrates including growth factors and their cognate receptors, adhesion molecules and chemokines^3^. For decades, MMPs have been studied in the context of cardiovascular diseases, rheumatoid arthritis and neurological disorders. MMPs are also of prime interest in cancer since most members of the MMP family have been found to be dysregulated in human cancers^4^.

MMPs are essential for key cellular processes from extracellular matrix regulation to cell migration and invasion to cell proliferation and apoptosis. Therefore, they are important in adults for wound healing, angiogenesis, tissues homeostasis and immunity. In addition, MMPs are expressed during embryogenesis and are involved throughout development^5^. Indeed, about half of all MMP family members are expressed during embryogenesis (see Table1).

**Table1.**
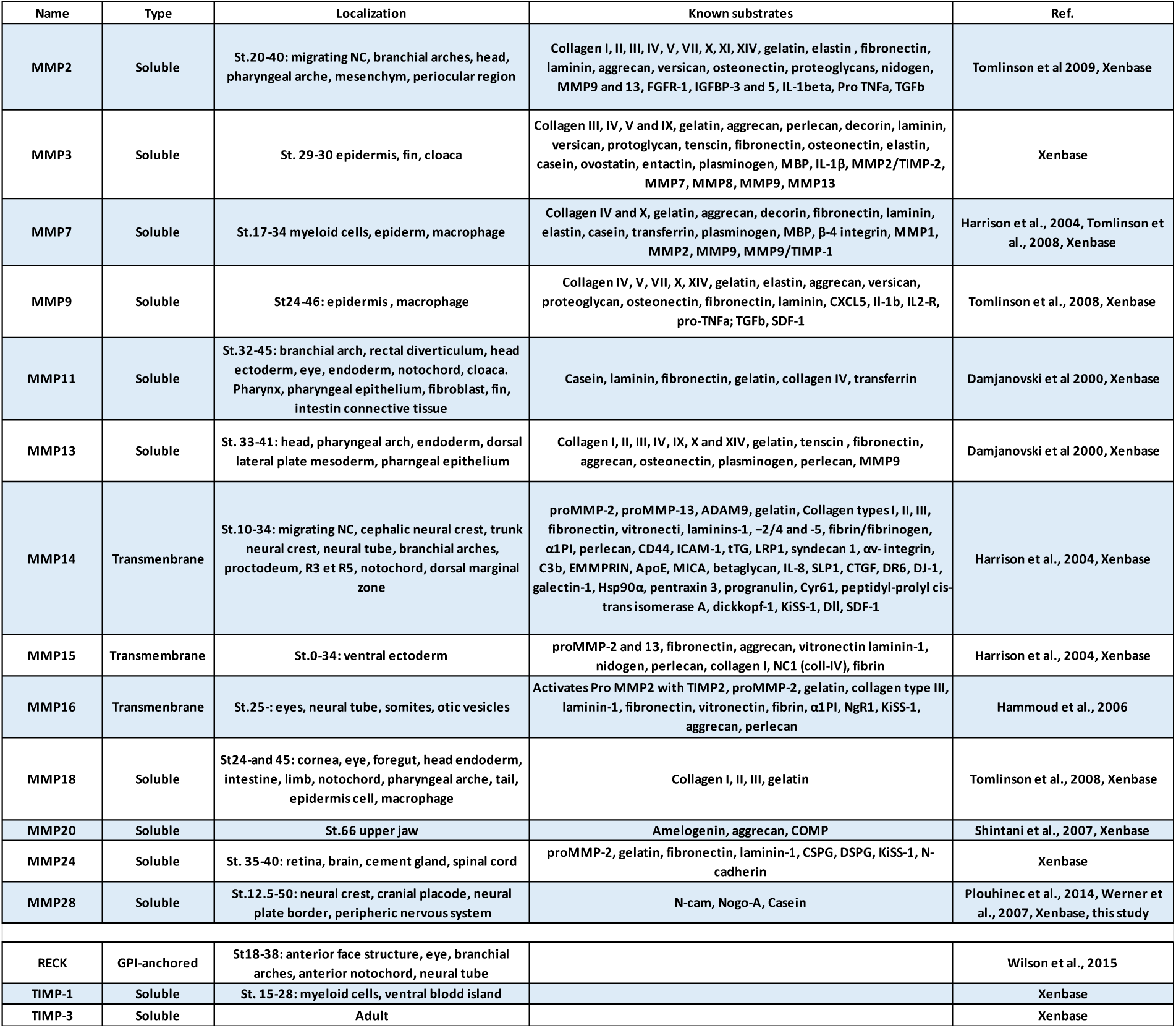
Developmental expression and putative substrates of matrix metalloproteinases identified in *Xenopus laevis*

MMP28 was first identified in human and is the last member of the family to be discovered^6,7^. MMP28 is a soluble MMP with endopeptidase activity and the ability to degrade casein, N-CAM and Nogo-A proteins^8^. MMP28 is involved in nerve repair and wound healing, immune system maturation, cell migration and invasion^9,10^. MMP28 is expressed during mouse and human embryogenesis^7,8,11^. In *Xenopus*, MMP28 was first identified in a screen for genes expressed in the neural crest (NC)^12^. NC cells arise from the neural plate border alongside the cranial placodes. They give rise to a wide variety of derivatives including smooth muscles, melanocytes, craniofacial bones and cartilages, and together with placode cells all the peripheral nervous system of the head^13^. So far MMP2, 11, 13, 14 and 16 were described as being expressed by the NC or by tissues surrounding the NC (see Table1). However, a detailed expression profile of MMP28 during *Xenopus* development is still lacking.

In this study we used qPCR analyses, single and double *in situ* hybridization (ISH) to assess the expression of MMP28 during *Xenopus laevis* embryogenesis. We found that MMP28 has a well-defined expression domain in the pre-placodal region at early neurula stage. During NC cells migration, MMP28 is then detected in some placodes and NC subpopulations. Given that NC and placodes interact to form the cephalic peripheral nervous system^13^ and that MMP28 is secreted, we propose that MMP28 may play a role in mediating these interactions to direct cranial morphogenesis.

## Materials and Methods

### *Xenopus* manipulation and in vitro fertilization

All experiments were performed in accordance with the guidelines of the Guide for the Care and Use of Laboratory Animals of the National Institutes of Health, and were approved by the Institutional Animal Care and Use Committee of New York University (animal protocol #IA16-00052). Female *Xenopus laevis* were injected with 500 to 1000 Units of chorionic gonadotrophin and kept overnight at 18°C. For fertilization, a suspension of minced testis was added to the oocytes collected in a petri dish in 0.1X Normal Amphibian Medium (NAM): NaCl (110mM), KCl (2mM), Ca(CO_3_)_2_ (1mM), MgSO_4_ (1mM), EDTA (0.1mM), NaHCO_3_ (1mM), Sodium Phosphate (2mM). Embryos were staged according to Nieuwkoop and Faber, 1967.

### qPCR

Embryos were collected by quick-freeze into liquid nitrogen. Total RNAs from batch of 3-5 embryos were extracted with the Rneasy® Micro Kit (Qiagen, Valencia CA). Relative quantitative PCR was performed on a QuantStudio 3 Real-Time PCR System (Applied Biosystems, Foster City, CA) using Power SYBR™ Green RNA-to-C_T_ 1-Step Kit according to manufacturer instructions and the following primers: Twist1qPCR_fdw: 5’-CGACTTTCTCTGCCAGGTCT-3’, Twist1qPCR_rev: 5’-TCCACACGGAGAAGGCATAG-3’, MMP28-E7_fdw: 5’-TGCAGTGGTATCGGGTTTAG-3’, MMP28-E8_rev: 5’-AAAGTGCAGTGTCAGGACGA-3’, Sox10_fdw: 5’-CTGTGAACACAGCATGCAAA-3’ Sox10_rev: 5’-TGGCCAACTGACCATGTAAA-3’, Six1_fdw: 5’-CTGGAGAGCCACCAGTTCTC-3’, Six1_rev: 5’-AGTGGTCTCCCCCTCAGTTT-3’ ODC_fdw: 5’-ACATGGCATTCTCCCTGAAG-3’, ODC_rev: 5’-TGGTCCCAAGGCTAAAGTTG-3’. All primer pairs were validated using the standard curve method. Data were normalized to ODC using the ΔCt method^14^.

### *In situ* hybridization and histology

Embryos were fixed for 1h at room temperature in MEMFA (0.1 M MOPS pH 7.42 mM EGTA pH 7.0, 1 mM MgSO_4_, 3.7 % (wt:vol) Formaldehyde) and store in 100% methanol. Embryos were then rehydrated in methanol solutions of decreasing concentrations, and processed for single or double ISH as previously described^15^. The following probes were used: Xl-Snai2^16^, Xl-MMP28, Xl-Sox10^17^, Xl-Six1^18^, Xl-Foxi4.1^19^, Xl-Sox2^20^ and Xl-Dmrta1^21^. For histology, stained embryos were embedded in Paraplast+, sectioned (12 µm) on an Olympus rotary microtome, counter stained with Eosin and mounted in Permount.

## Results

*Xenopus* has two homologous MMP28 genes on chromosome 2L (MMP28.L) and 2S (MMP28.S), which are composed of 8 exons each and encode a 1977 bp and 1916 bp mRNAs, respectively. Both mRNA sequences share 84.71% identity, and 92.9% identity for the open reading frame. MMP28.L and MMP28.S code for proteins of 496 and 497 amino acids, respectively, with a predicted molecular weight of 57kDa. Both protein sequences share 91.15% identity and 97.58% similarity. As the sequences share high identity, we designed primers and probes for ISH that detect both MMP28.L and MMP28.S.

We assessed the temporal expression of MMP28 by relative qPCR at different stages of *Xenopus* development from stage 4 (8-cell stage) to tailbud stage (Figure 1, blue bars), and compared its expression profile to that of Twist, a marker of cephalic NC^22^ (Figure 1, light grey bars) and Six1, a marker for cranial placodes^23^ (Figure 1, black bars). MMP28 is maternally expressed (stage 4) although at relatively low levels, similar to Six1, while Twist does not appear to have a maternal contribution. After the mid-blastula transition the expression of embryonic genes is initiated. MMP28 expression levels remain relatively low at stage 11 and stage 12.5 while both Twist and Six1 expressions increase significantly during that period until stage 15 where they both reach a plateau of expression. MMP28 shows a marked increased expression at stage 14, which is maintained until stage 18. Later in development, MMP28 expression progressively increases up to stage 27, the last stage examined in this study. Twist reaches its higher expression level at stage 18 without any further increase later in development. Six1 expression decreases significantly at stage 18 and at later stage progressively increases similarly to MMP28. These data indicate that MMP28 expression is strongly upregulated at the neurula stage and the escalation of Twist and Six1 expression precedes that of MMP28.

**Figure 1.**
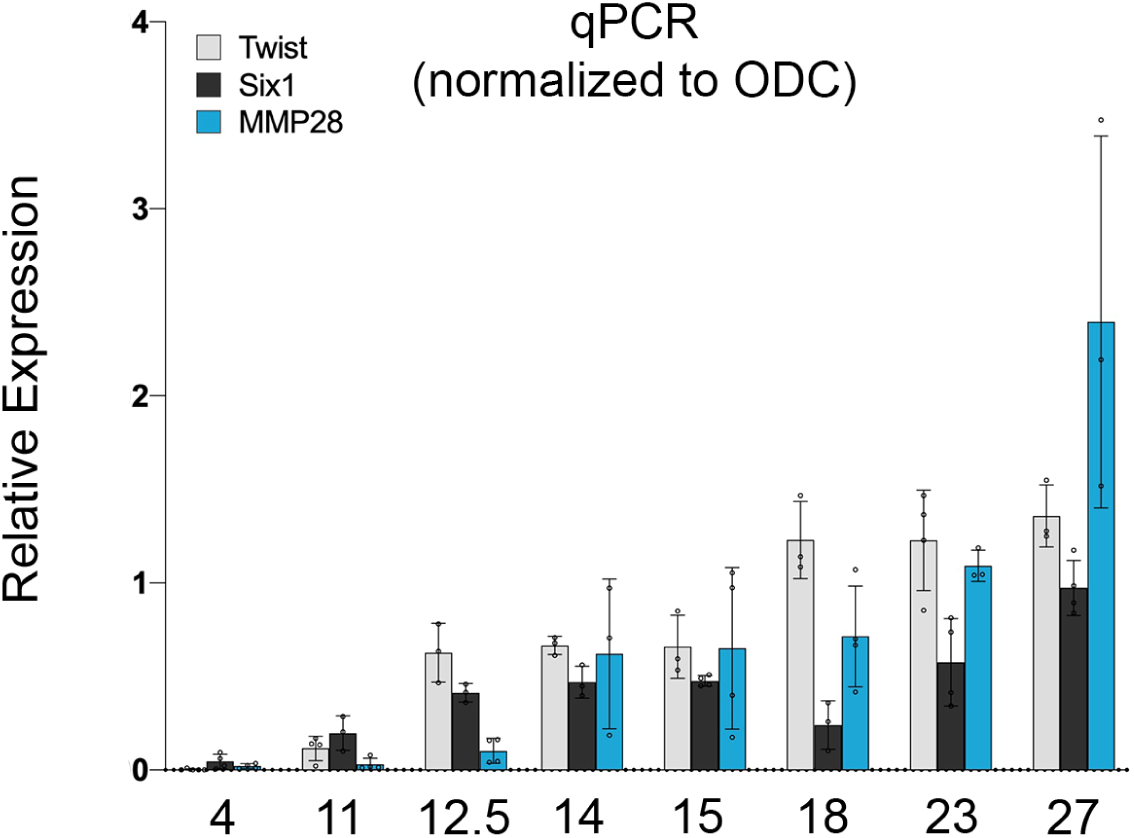
Relative expression of Twist (light grey), Six1 (black), MMP28 (blue) during *Xenopus* embryogenesis from stage 4 to stage 27. Results are expressed using ΔCt method with the house-keeping gene ODC as reference. For representation purposes, relative expression values results were multiplied by 1000.

We next performed ISH to characterize the spatial expression of MMP28 during *Xenopus* development. MMP28 is not detected at gastrula stage (Figure 2 A, B). MMP28 transcripts first accumulate at detectable levels at the end of gastrulation (NF stage 12.5), in the form of a thin horseshoe shape domain at the anterior part of the embryo, which represents the neural plate border (Figure 2 E), the region that gives rise to both cranial placodes and NC cells. A control sense probe confirmed the specificity of this expression (Figure 2 D). At stage 14/15, MMP28 expression is more defined into two bilateral domains on either side of the neural plate (Figure 2 F-H). This staining strongly suggests a placodal expression since the neural fold is devoid of staining. At this stage, MMP28 is also detected in a more discrete domain along the border of the neural plate (Figure 2 H and I’, arrowhead). To substantiate these observations, stage 15 embryos were sectioned to visualize MMP28 expression in more detail (Figure 2 H, H’, H’’ and H’’’).

**Figure 2.**
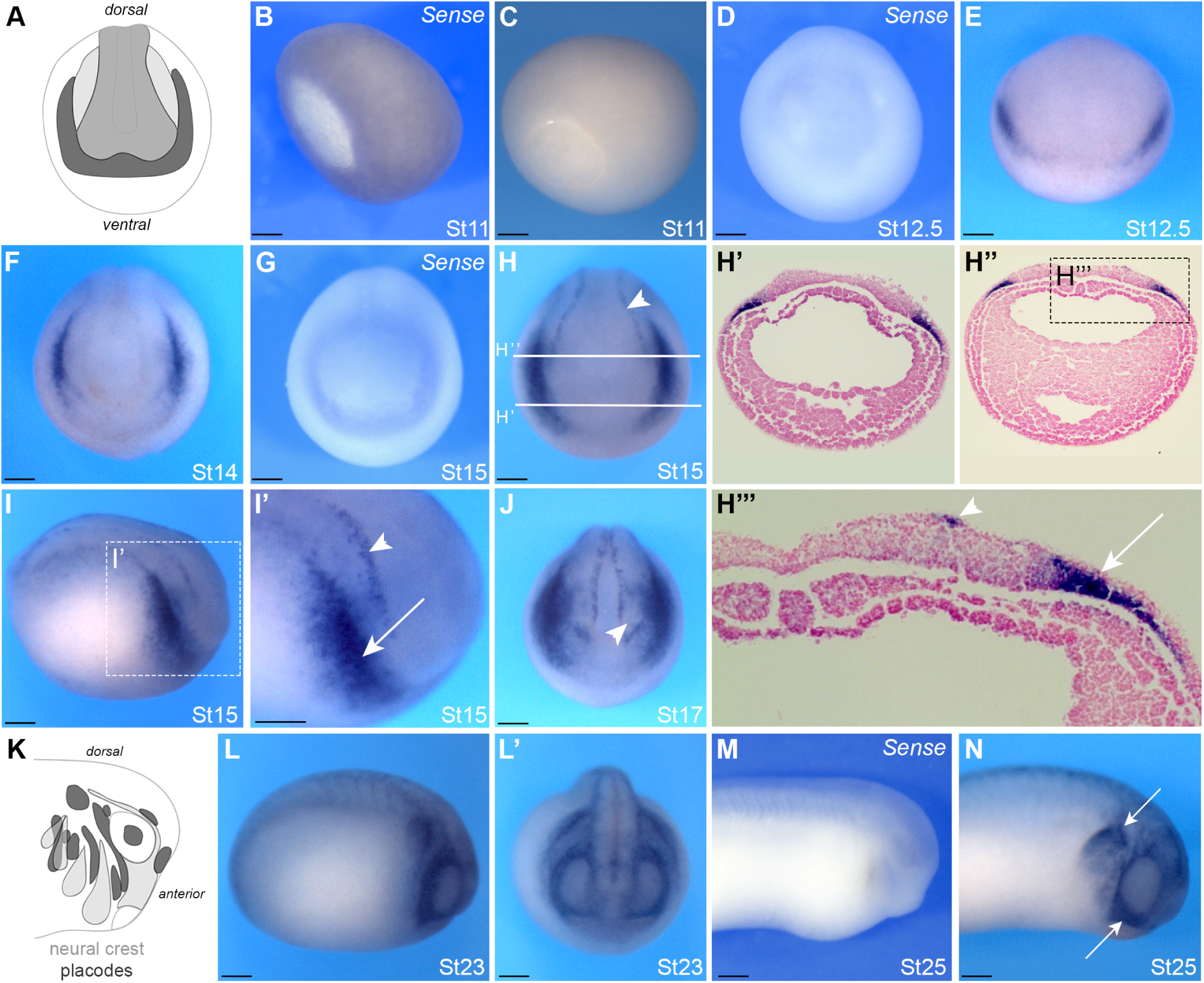
Developmental expression of MMP28 by In Situ Hybridization (ISH). ISH for MMP28 in lateral view (B, C, I, I’, L, M, N), anterior view (D-E, J, L’), dorsal view (F-H) and transversally sectioned embryos (H’ and H’’). A and K: schematic representation of placodes (dark grey), neural crest (light grey) and neural plate (grey). (B-C) At St 11 MMP28 is not detected. (D-E) MMP28 is first detected at the anterior neural plate border at St 12.5. (F-J) Neurula stage embryos show expression in the lateral placodes (arrow) and medial NC (arrow-head). (H’-H’’’) Transversal sections as indicated in H, placodes (arrow), medial NC (arrowhead). (L-L’) Expression in the epibranchial placodes. (M-N) At tailbud stage MMP28 expression is detected in the first NC stream (arrows). The embryonic stage is indicated in the lower right corner of each panel. When MMP28 sense probe is used (Sense) it is indicated in the upper right corner of the panel. Scale bar: 0.25mm

MMP28 expression is localized in the deep ectoderm layers consistent with cranial placodes expression (Figure 2 H’’’ and I’, arrow). By contrast, the thin line of expression along the neural plate is localized in the most superficial ectoderm layer (Figure 2 I’ and H’’’, arrowhead), and correspond the medial NC^24^. At stage 17 the expression is broader and outlines the prospective eyes (Figure 2 J, arrowhead). At stage 23, during NC cells migration, MMP28 expression seems to be restricted to discrete domains around the eyes and in territories posterior to that region, likely to correspond to the epibranchial placodes (Figure 2 L, L’). At stage 25, strong MMP28 expression is detected in the first stream of NC as well as what appears to be the third and fourth NC streams (Figure 2 N, arrows). The specificity of MMP28 expression was confirmed using a control sense probe (Figure 2 M).

In order to identify more precisely which tissues express MMP28, we performed ISH at stages 15 and 25 for MMP28 and genes expressed in similar regions of the ectoderm, including Sox2 (neural plate and cranial placodes), Snai2/Slug (NC), Dmrt1a (olfactory placodes), Foxi4.1 (epibranchial placodes) and Sox10 (migrating NC). The probes were used alone or in combination as indicated (Figure 3). Spatially there is no overlap between MMP28 and Snai2, as Snai2 expression domain (Figure 3 A, A’) is medial to that of MMP28 (Figure 3 G, G’ and B, B’). Sox2 is expressed into two regions, the neural plate (Figure 3 C, arrow) and lateral placodes (Figure 3 C and C’, arrowhead). MMP28-positive territory overlaps exclusively with Sox2 placodal expression domain (Figure 3 D, D’). Since the placodal domain gives rise to a broad range of different placodes, we also compared MMP28 to Dmrta1, a gene restricted to the most anterior placodal region, the prospective olfactory placodes (Figure 3 E, E’). We found that MMP28 and Dmrta1 are expressed in adjacent but non-overlapping domains (Figure 3 F, F’). At stage 25, Foxi4.1 is strongly expressed in the epibranchial placodes and sensory layer of the ectoderm (Figure 3 J, arrowheads) but not in the NC streams (Figure 3 J, arrows). In contrast Sox10 is expressed strongly in all NC streams (Figure 3, arrows) and excluded from the branchial ectoderm. This histological analysis indicates that, contrary to what whole-mount ISH staining suggested, MMP28 is not expressed in the posterior NC streams (Figure 3 L, arrows) but shows clear expression in the epibranchial placodes and sensory layer of the ectoderm (Figure 3 L, arrowheads).

**Figure 3.**
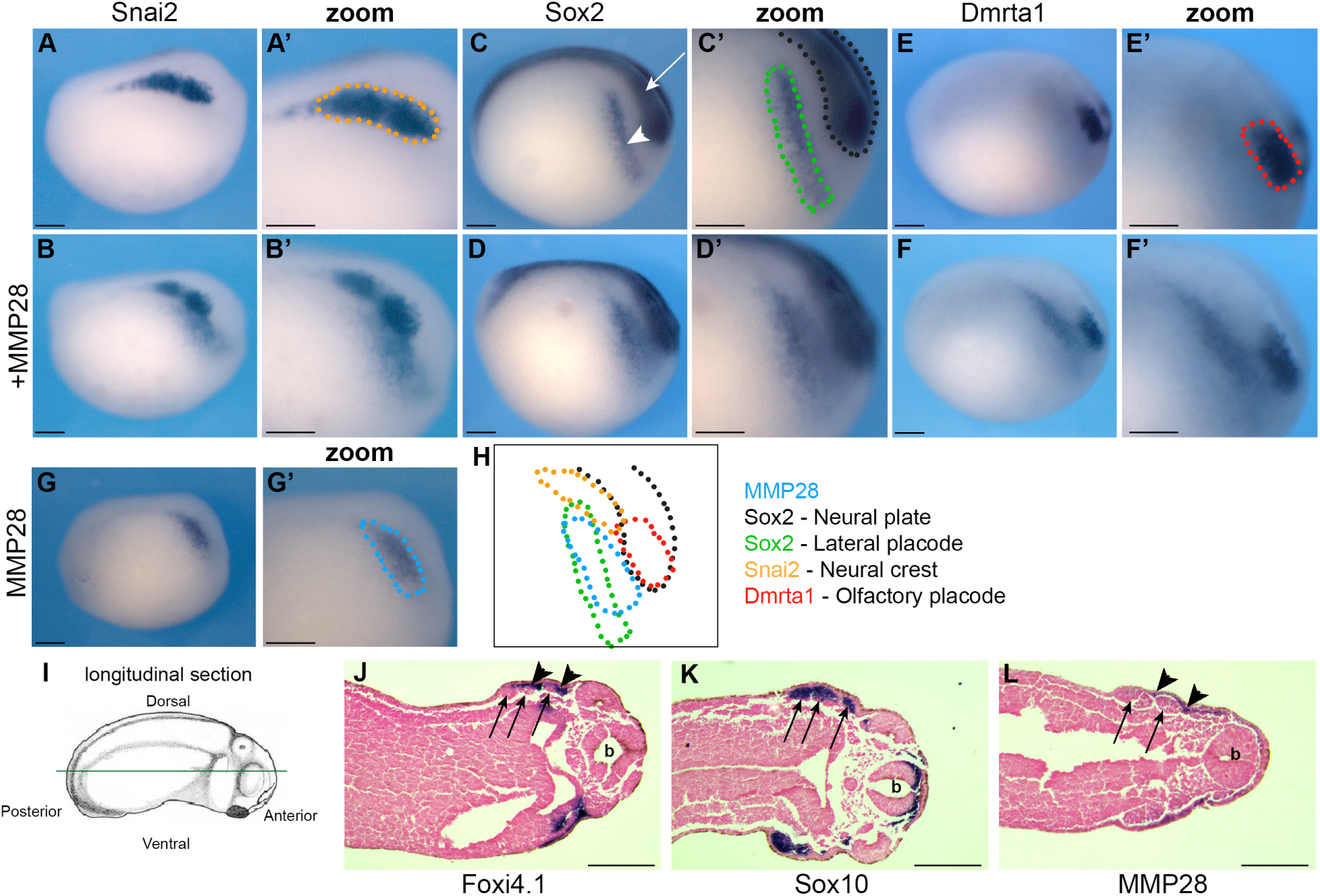
Comparative analysis of MMP28, Snai2, Sox2, Dmrta1, Foxi4.1 and Sox10 expression by single and double ISH. A-H: *Xenopus* embryos at stage 15 (lateral view, anterior to right, dorsal to top) after single ISH for Snai2 (A-A’), Sox2 (C-C’), Dmrta1 (E-E’) and MMP28 (G-G’), or double ISH for MMP28/Snai2 (B-B’), MMP28/Sox2 (D-D’) and MMP28/Dmrta1 (F-F’). Summary of the expression domains of these genes (H). Doted lines represent expression territory of MMP28 (blue), Snai2 (orange), Sox2 (black, neural plate; green, placodes) and Dmrta1 (red). MMP28 (blue) overlaps with the placodal domain of Sox2 (green). I-L: *Xenopus* embryos at stage 25 (anterior to the right, posterior to the left). Diagram illustrating the plane of section (green line; I). Single ISH for Foxi4.1 (J), Sox10 (K) and MMP28 (L). Arrows show neural crest streams, arrowheads show epibranchial placodes, b: brain. Scale bar 0.25mm.

Our results indicate that, at neurula stage, MMP28 is not expressed in the NC but is confined to the cranial placodes territory overlapping with Sox2 (Figure 3 H) and that its expression is maintained in the epibranchial placodes and sensory layer of the ectoderm at tailbud stage (Figure 3 L).

## Discussion

Here we report the developmental expression of MMP28, a member of the matrix metalloproteinases family, in *Xenopus*. Our results showed that MMP28 is expressed maternally even though faintly and is upregulated at the end of gastrulation (St. 12.5) at which point it becomes detectable by ISH. We observed that MMP28 is uniquely expressed at the neurula stage in the pre-placodal region adjacent to the prospective NC territory.

We showed that MMP28 expression is limited to the neural plate border at early neurula stage, then to the lateral placodes. This placodal expression is sustained in the epibranchial placodes at tailbud stage. In addition, MMP28 expression is detected along the migratory pathway of the first NC stream. As indicated earlier, MMP28 was first isolated in a screen for genes activated in the NC tissue at neurula stage^25^. However, because the microarray-based approach was performed on manually dissected embryonic tissues at different stages, potential contamination by surrounding tissues including neural plate, neural folds, neural crest or lateral ectoderm cannot be excluded. Indeed, at neurula stage, our double ISH show that MMP28-positive signal does not overlap with the NC marker Snai2/Slug, but colocalized with the lateral placodal domain of Sox2. While we have detected expression in the medial NC overall, MMP28 expression is primarily confined to the cranial placodes at early neurula stages.

The fact that MMP28 is strongly expressed in placodes is very interesting and unique. MMP1, 2, 3, 7, 9, 11, 13, 14, 15, 18, 20 and 24 are expressed during *Xenopus* embryogenesis (Table1). Almost half of them are expressed by the NC or surrounding tissues and their derivatives, but none of them has been described in placode cells at the stages studied here. MMP28 expression profile overtime is comparable to that of Six1. However, spatially MMP28-positive territory only represents a subdomain of Six1^23^. Notably, MMP28 was not detected in the most anterior placodal domain giving rise to olfactory placodes. By contrast, MMP28 expression is maintained in the epibranchial placodes that originate from the lateral placodes (Sox2-positive territory) and later is detected in between migratory NC streams.

NC cells and cranial placodes cooperate to form the cephalic peripheral nervous system. NC cells give rise to glial cells while neurons have a dual origin coming from either NC, placodes or both depending on the cranial ganglion of interest^26^. Interestingly, NC and placodes interaction involved in peripheral nervous system patterning starts during NC migration. Indeed, in *Xenopus*, placodal cells are the source of Stromal cell-derived factor 1 (Sdf1/Cxcl12) a key regulator of NC cell migration^27,28^. Sdf1 promotes NC cell motility, which in the context of the constraints of the developing head leads to migration of NC cells towards the placodes. However, physical interactions between the two cell types leads to a repulsion. This short range repulsion mid-range attraction system, coined “chase-and-run”, leads to a sustained directional migration of both NC and placode cells towards the ventral regions of the face^29^. Placodal cells are less motile than NC cells and are progressively pushed aside while NC cells migrate ventrally. This leads to a pattern of accumulation of placode cells in between NC streams^29,30^. Therefore, interfering with NC migration leads to defects in placode patterning and this relationship is conserved in *Xenopus* and zebrafish^29,31^. The expression of MMP28, a secreted MMP, in the placodes prior to the onset for NC migration raises the possibility that this enzyme might be required in normal NC-placode interactions.

## Conclusion

Overall our data indicate that the onset of MMP28 expression in placodes occurs after the NC and pre-placodal domains have been specified at the neural plate border. MMP28 expression is excluded from the anterior-most placodes and persists in epibranchial placodes strongly suggesting a role for this molecule in NC-placodes interactions at early stages of the cranial peripheral nervous system formation.

## Acknowledgment

The authors would like to thank Ms. Allison Williams for excellent technical assistance.

